# Mutations in the catalytic domain of human β-cardiac myosin that cause early onset hypertrophic cardiomyopathy significantly increase the fundamental parameters that determine ensemble force and velocity

**DOI:** 10.1101/067066

**Authors:** Arjun S. Adhikari, Kristina B. Kooiker, Chao Liu, Saswata S. Sarkar, Daniel Bernstein, James A. Spudich, Kathleen M. Ruppel

## Abstract

Hypertrophic cardiomyopathy (HCM) is a heritable cardiovascular disorder that affects 1 in 500 people. In infants it can be particularly severe and it is the leading cause of sudden cardiac death in pediatric populations. A high percentage of HCM is attributed to mutations in β-cardiac myosin, the motor protein that powers ventricular contraction. This study reports how two mutations that cause early-onset HCM, D239N and H251N, affect the mechanical output of human β-cardiac myosin at the molecular level. We observe extremely large increases (25% – 95%) in the actin gliding velocity, single molecule intrinsic force, and ATPase activity of the two mutant myosin motors compared to wild type myosin. In contrast to previous studies of HCM-causing mutations in human β-cardiac myosin, these mutations were striking in that they caused changes in biomechanical parameters that were both greater in magnitude and more uniformly consistent with a hyper-contractile phenotype. In addition, S1-S2 binding studies revealed a significant decrease in affinity of the H251N motor for S2, suggesting that this mutation may further increase hyper-contractility by releasing active motors from a sequestered state. This report shows, for the first time, a clear and significant gain in function for all tested molecular biomechanical parameters due to HCM mutations in human β-cardiac myosin.

## Introduction

Hypertrophic cardiomyopathy (HCM) is a heritable cardiovascular disorder characterized by abnormal thickening of the left ventricular walls (*1*), preserved or increased systolic function and reduced diastolic function (*2*). HCM is typically diagnosed in late adolescence or adulthood, and is the leading cause of sudden cardiac death in those under the age of 35 (*3*). Disease presentation in infancy or childhood is associated with poorer outcomes; rates of death/cardiac transplantation approach 20% (*4*).

HCM is most commonly caused by mutations in genes encoding sarcomeric proteins, principally those encoding β-cardiac myosin (MYH7) and cardiac myosin binding protein-C (MYBP3) (*5–9*). Despite the identification of the genetic basis of this disease, relatively little is known about the molecular mechanisms by which these mutations result in the HCM disease phenotype. It has been hypothesized that mutations in β-cardiac myosin, the mechanoenzyme that drives ventricular contraction, cause HCM by affecting the power output of the myosin motor (*10*). Power output is the product of force and velocity, and so the effects of HCM-causing mutations on myosin power output at the molecular level can be determined by assaying the velocity of actin filament gliding on ensembles of mutant myosin motors and the force produced by those ensembles. Ensemble force can be estimated by determining the key parameters that contribute F_ensemble_ = F_intrinsic_ (ts/tc) Na, where F_intrinsic_ is the intrinsic force of the motor, t_s_ is the time of the ATPase cycle that myosin is tightly bound to actin, t_c_ is the total cycle time of the actin-activated myosin ATPase, t_s_/t_c_ is the duty ratio, and N_a_ is the number of myosin heads that are functionally accessible for interaction with actin in the sarcomere.

Previous studies of the effects of HCM mutations on these biomechanical parameters have shown highly variable results (**Table 1**). Studies on R403Q, one of the most extensively investigated HCM mutations, performed with mouse α-cardiac myosin have shown 30-120% increases in ATPase activity, 10-30% increases in velocity and a 50% increase in force (*11–13*). However, studies done on mouse β-cardiac myosin showed no change in the velocity and ATPase activity (*12*), while patient derived human β-cardiac myosin showed a decrease in force (*14*). For R453C, experiments done with mouse α-cardiac myosin showed no change in velocity (*11, 13*), either no change (*11*) or 45% decrease (*13*) in ATPase activity, and an 80% increase in force (*11*).

**Table 1.** Myosin biomechanical parameters for myosins carrying various HCM mutations relative to WT myosin of the corresponding species. Increases over WT are in blue, decreases in red. NC = no change. Not determined (-).

| HCM mutation | Myosin | Velocity | ATPase | Force |
| --- | --- | --- | --- | --- |
| R453C(13) | Mouse α-cardiac | NC | 0.6 | - |
| R453C(11) | Mouse α-cardiac | NC | NC | 1.8 |
| R403Q(11) | Mouse α-cardiac | 1.3 | 2.2 | 1.5 |
| R403Q(12) | Mouse α-cardiac | 1.3 | 1.3 | - |
| R403Q(11) | Mouse α-cardiac | 1.1 | 1.4 | - |
| R403Q(12) | Mouse β-cardiac | NC | NC | - |
| R403Q(43) | Human β-cardiac<br>(human tissue) | 1.3 | - | - |
| R403Q(14) | Human β-cardiac<br>(human tissue) | - | - | 0.6 |
| R453C(19) | Human β-cardiac sS1 | 0.8 | 0.7 | 1.5 |
| R403Q(18) | Human β-cardiac sS1 | 1.1 | 1.2 | 0.8 |
| R719W(20) | Human β-cardiac sS1 | 1.1 | NC | 0.8 |
| R723G(20) | Human β-cardiac sS1 | 1.1 | NC | 0.7 |
| G741R(20) | Human β-cardiac sS1 | NC | NC | NC |
| H251N (this manuscript) | Human β-cardiac sS1 | 1.4 | 1.3 | 1.4 |
| D239N (this manuscript) | Human β-cardiac sS1 | 1.9 | 1.5 | 1.2 |

Studies to define the functional consequences of these and other β-cardiac myosin mutations have been hampered by the lack of available expressed and purified human cardiac myosin, but this has been solved by the establishment of a mouse myoblast expression system (*15–17*). Thus, highly-purified recombinant human β-cardiac myosin short S1 (sS1) motor domains have been examined for the effects of several HCM mutant forms of human β-cardiac sS1(*18–20*). However, the results using these recombinant human motors have been variable, with no clear pattern emerging (*18, 19*). Most mutations show small differences in one or more parameters compared to WT myosin, and typically a hyper-contractile change in one parameter is accompanied by a hypo-contractile change in another. For example, R403Q led to a gain in velocity (10%) and ATPase activity (25%), but a decrease (15%) in force (*18*). R453C caused a decrease in velocity (25%) and ATPase activity (30%), but a large increase in force (50%) (*19*). Recently, Kawana et al. (*20*) studied three converter domain mutations, and found very subtle changes. For the converter mutations R719W and R723G, there was a ~5-10% increase in velocity, no change in ATPase activity, and 20-30% decrease in force, respectively, while G741R showed no significant change in velocity, ATPase activity, or force (*20*). It is unclear whether these small changes reflect the fact that mutations that give rise to HCM cause relatively small perturbations in myosin biomechanics or rather that current assays do not accurately measure the effects of HCM mutations on β-cardiac myosin’s force-velocity relationship.

In order to address this question, we studied the effects of mutations that cause severe, early onset HCM. Although the causative mutation is present at birth, most individuals with HCM do not manifest evidence of the disease until late adolescence or adulthood. However, in a subset of individuals, symptomatic HCM can present in infancy or childhood (pediatric HCM incidence ~5 per million) (*4*). Disease progression can be swift, with significant morbidity and mortality, especially in infants, who account for the vast majority of pediatric HCM (*4*). Three quarters of all pediatric-onset HCM is classified as idiopathic (*4*). It is now known that 50-60% of idiopathic HCM is caused by mutations in genes encoding sarcomeric proteins, and roughly half of those are in β-cardiac myosin (*21, 22*). Recently, novel β-cardiac mutations have been found that cause severe disease in infants and children (*21, 22*). Some of the identified pediatric HCM mutations have not been reported, or reported only rarely, in adult patient populations (*23*). While the number of patients with these mutations is small, statistical analysis suggests that these mutations may be associated with an HCM subtype characterized by more severe, early onset disease (Knight and Leinwand, personal communication). We chose two early onset HCM mutations, H251N on the myosin mesa, a relatively flat surface of the catalytic domain that has been proposed to be a binding site for the myosin coil-coiled tail and myosin binding protein-C (*24, 25*), and D239N within the Switch-1 (nucleotide binding) domain (**Figure 1**), to test whether mutations associated with such a severe clinical phenotype would exhibit more significant, and consistently hyper-contractile, changes in myosin biomechanical parameters. We utilized assays for actin-activated myosin ATPase activity (*26*), in vitro motility (*27*), loaded in vitro motility (*28*), and single molecule intrinsic force (F_intrinsic_) by optical tweezers force spectroscopy (*29*) to elucidate the effect of each mutation on biochemical cross-bridge cycling parameters. We also analyzed binding between myosin S1 and S2 domains (*30*) to examine the hypothesis that HCM mutations in the myosin mesa disrupt a sequestered state of myosin heads in the sarcomere that limit the number of functionally accessible heads for interaction with actin (*24*). Together, we show that these mutations exhibit clear, significant gain of function, unlike many recently studied HCM-causing mutations in human β-cardiac myosin.

**Figure 1.**
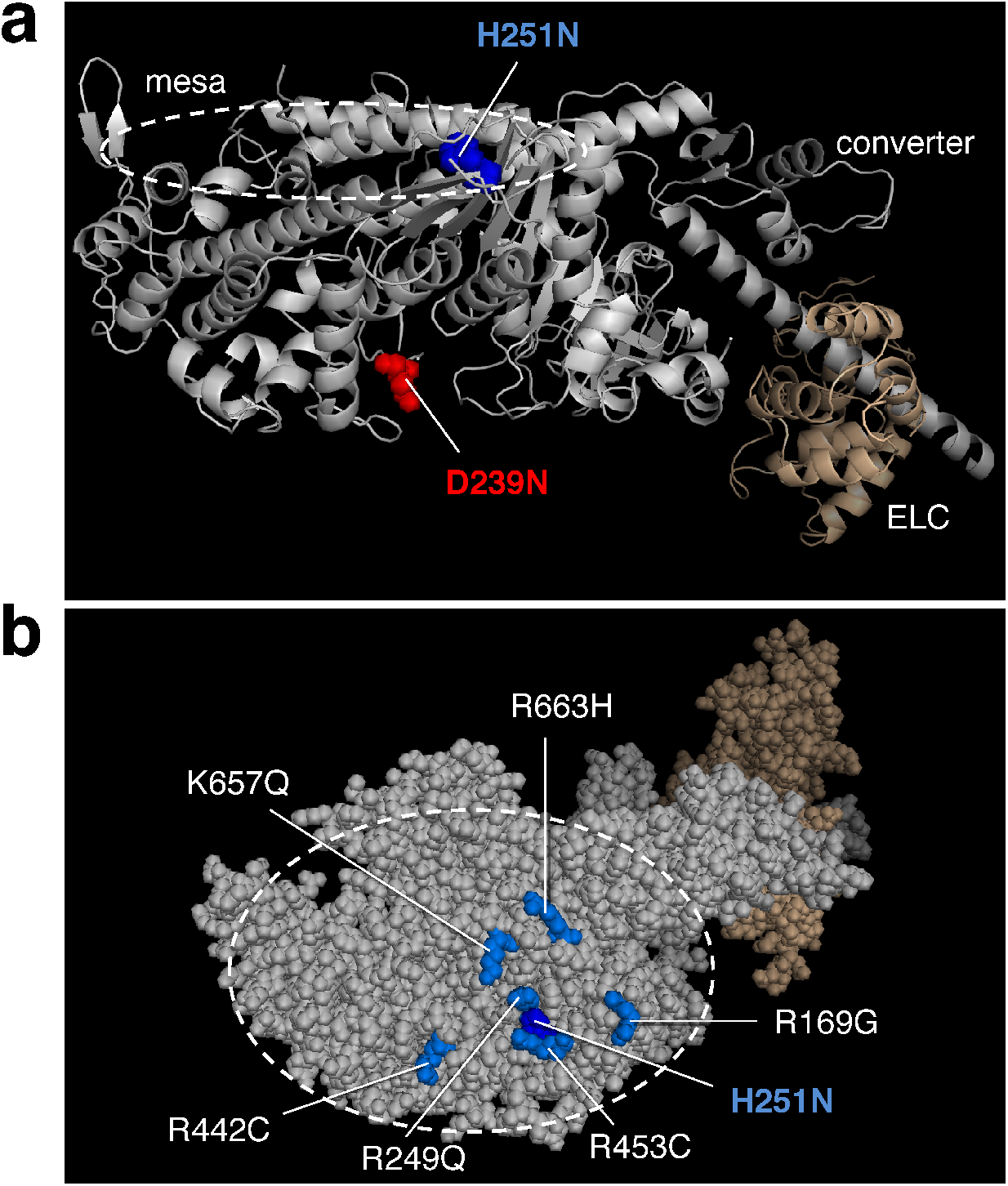
Structure of a homology-modeled human β-cardiac sS1 domain for two pediatric early onset HCM mutant forms of the protein. (a) Structure of homology-modeled (see Methods) human β-cardiac sS1, which contains residues 1-808 of the MyHC (grey) and the ELC (brown). The positions of the HCM mutations H251N (blue) and D239N (blue) are shown. (b) Sphere model of sS1 shown in (a) rotated ~90° about the horizontal axis toward the reader, viewing the mesa from the top. The position of the mutation H251N on the mesa is seen in the middle of a cluster of other positively charged residues, all of which cause HCM (*23*).

## Results

### H251N and D239N have significantly increased actin-activated ATPase activity compared to WT

To assess the effects of the mutations on total cycle time (t_c_), we measured the maximal ATPase activity of the actin-myosin complex (k_cat_) for WT and each mutant (t_c_ = 1/k_cat_). We found that the k_cat_ for H251N (5.6 ± 0.4 s^-1^, 8 replicates from 3 sS1 preparations) and for D239N (6.7 ± 0.5 s^-1^, 6 replicates from 3 sS1 preparations) were significantly increased compared to WT sS1 (4.5 ± 0.4 s^-1^,12 replicates, 5 sS1 preparations) (**Figure 2** and **Table 2**). To our knowledge, these increases in catalytic rates (24% for H251N and 50% for D239N) are the largest observed for any mutation assayed in the human β-cardiac myosin backbone, and thus the largest decreases in t_c_. There were no changes in the values of K_M_ for either of the human β-cardiac mutant myosins as compared to WT (**Table 2**).

**Figure 2.**
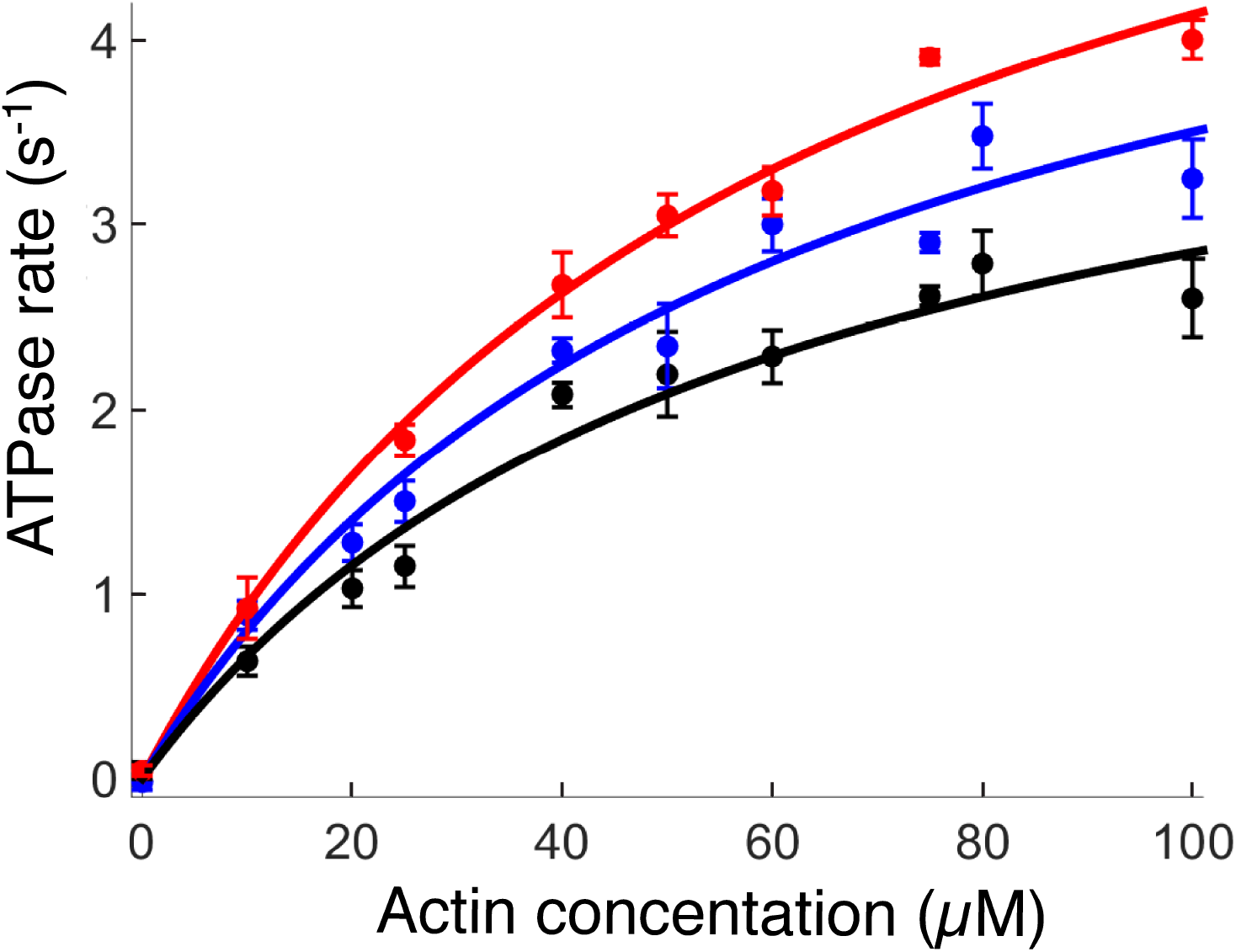
Actin-activated ATPase activity for two pediatric early onset HCM mutations. The actin-activated ATPase rate of sS1 is fitted to a Michaelis-Menten curve. WT is black, H251N blue and D239N red throughout all figures. The error bars are the standard error on the mean (S.E.M.)

**Table 2.**
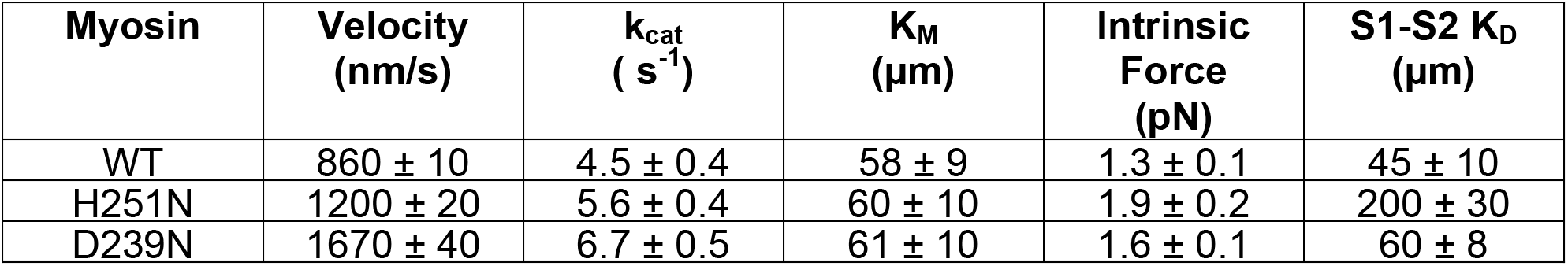
Biomechanical parameters for WT and mutant sS1s. Velocity was measured using the in vitro motility assay, k_cat_ and K_M_ were measured using an ATPase assay, and single molecule intrinsic force using optical tweezers. Each measurement was done with at least two separate preparations of sS1, with at least three independent runs each. The error reported is the standard error of the mean.

| Myosin | Velocity<br>(nm/s) | $k_{\text{cat}}$<br>($\text{s}^{-1}$) | $K_{\text{M}}$<br>($\mu\text{m}$) | Intrinsic<br>Force<br>(pN) | S1-S2 $K_{\text{D}}$<br>($\mu\text{m}$) |
| --- | --- | --- | --- | --- | --- |
| WT | $860 \pm 10$ | $4.5 \pm 0.4$ | $58 \pm 9$ | $1.3 \pm 0.1$ | $45 \pm 10$ |
| H251N | $1200 \pm 20$ | $5.6 \pm 0.4$ | $60 \pm 10$ | $1.9 \pm 0.2$ | $200 \pm 30$ |
| D239N | $1670 \pm 40$ | $6.7 \pm 0.5$ | $61 \pm 10$ | $1.6 \pm 0.1$ | $60 \pm 8$ |

### Single molecule optical tweezer force spectroscopy shows an increase in the intrinsic force of both H251N and D239N

To measure the effect of these mutations on the intrinsic force of human β-cardiac sS1, we used a dual beam optical trap, as described previously (*29*). Force histograms of individual molecules were used to calculate the mean force for each of the proteins (*18*) (**Figure 3a**). The WT sS1 single molecule mean force was measured as 1.3 ± 0.1 pN (2 preparations, 6 molecules). Compared to the WT sS1, both H251N (1.9 ± 0.2 pN; 2 preparations, 6 molecules) and D239N (1.6 ± 0.1 pN; 2 preparations, 6 molecules) showed appreciable increases in force (H251N = 46% and D239N = 23% increase, respectively) (**Figure 3a**). Additionally, we used a cumulative probability distribution function to independently assess the increase in contractile force at the single molecule level (*18*). For all the events measured with all the molecules, we calculated the cumulative probability distribution (*18*). From **Figure 3b** it is clear that at all cumulative probability values, the single molecule force of WT < D239N < H251 N.

**Figure 3.**
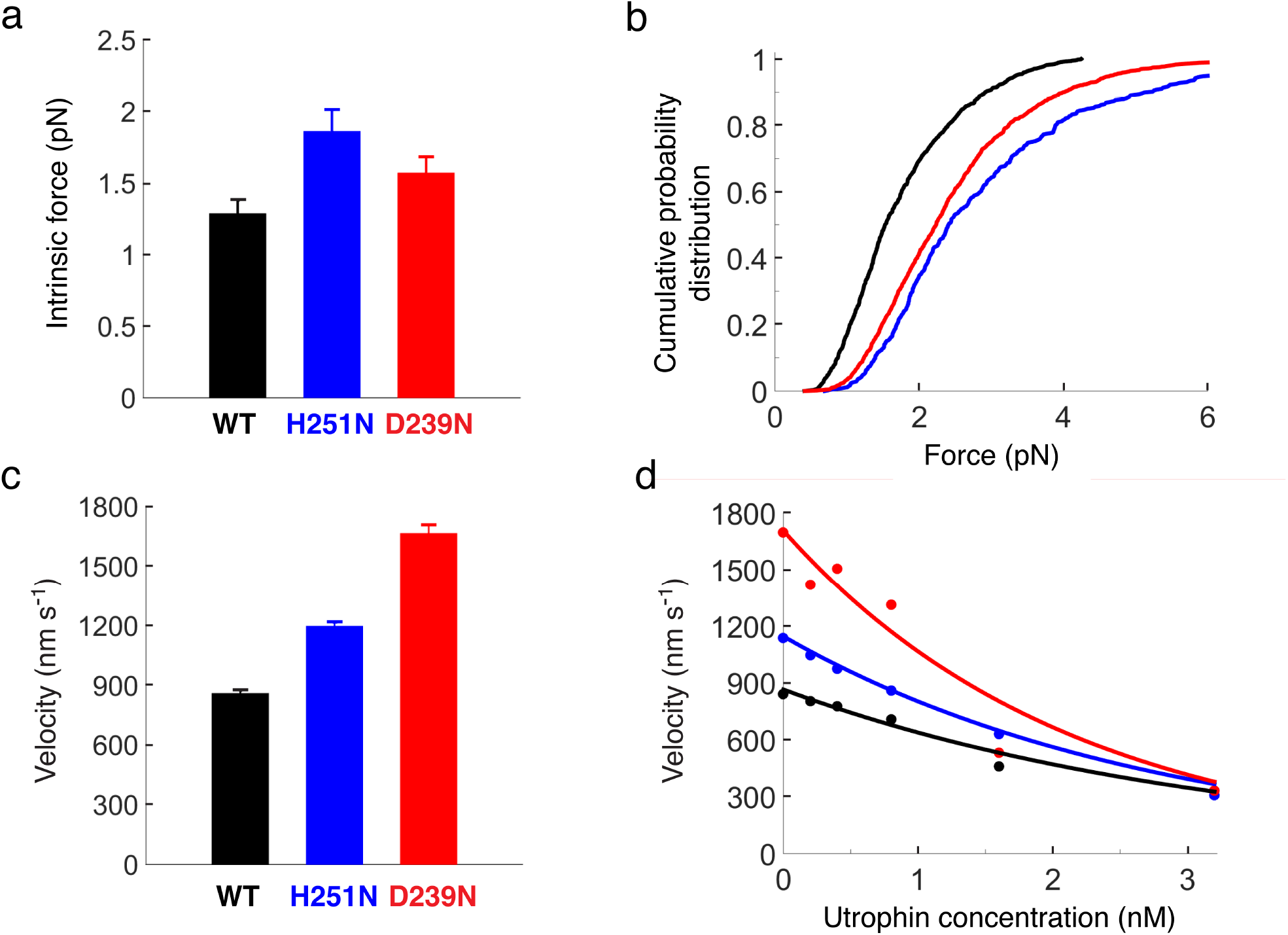
Single molecule optical tweezer and velocity measurements for H251N and D239N human β-cardiac sS1s. (**a**) Intrinsic force measure in a dual beam single molecule laser trap. (**b**) Force differences shown by a cumulative probability distribution. (**c**) Unloaded in vitro motility assay results. (**d**) Representative loaded in vitro motility results, using the actin binding protein utrophin as a load molecule (the fits serve as a guide to the eye). All error bars are S.E.M.

### H251N and D239N generate higher actin gliding velocities than WT in an *in vitro* motility assay

We next assessed the effect of the mutations on the velocity at which human β-cardiac sS1 propels actin filaments in an *in vitro* motility assay (*27, 31*). The average unloaded velocities of H251N (1200 ± 20 nm/s,10 replicates from 3 sS1 preparations) and D239N (1670 ± 40 nm/s, 6 replicates from 3 sS1 preparations) were significantly greater than that of WT sS1 (860 ± 10 nm/s, 25 replicates from 5 sS1 preparations) (**Figure 3c, Table 2**). As with the catalytic rate constants, these extremely large increases in functionality compared to the WT (D239N = 94% and H251N = 40%) are the largest changes observed for human β-cardiac myosin containing HCM-causing mutations.

### H251N and D239N both generate higher ensemble force than WT in a loaded *in vitro* motility assay

To gain further insight into how the biomechanical properties of myosin are altered due to these mutations, we utilized a loaded *in vitro* motility assay. Our loaded *in vitro* motility assay is an adaptation of the actin gliding assay, where the actin binding protein utrophin is introduced to act as a load molecule against which myosin works (*28*). Performing the motility assay at various utrophin conditions gives a load-velocity curve for the ensemble of myosin motors on the surface. (**Figure 3d**) (*28*). Examination of the curves for the human β-cardiac WT and mutant sS1s shows that at each applied load (utrophin concentration) the velocities of the mutant motors were higher than that of WT, suggesting a higher relative ensemble force and power output (power = force x velocity) for the two mutants.

### H251N sS1 binds to human cardiac proximal S2 with significantly decreased affinity compared to WT

Structural studies of several types of myosins, including cardiac myosin, have shown that the myosin S1 head domains fold back onto the proximal portion of the α-helical coiled-coil tail of myosin (*32–35*). It has been hypothesized that such folding back could sequester heads and thus alter the number of functionally accessible myosin heads for exerting force on actin (*24, 25, 32–35*). It has also been suggested that HCM-causing mutations may disrupt this sequestered state, releasing more heads to interact with actin and resulting in the hypercontractility phenotype observed clinically (*24, 25*). Indeed, Nag et al. (*25*) have shown that the myosin mesa mutation R453C, which is located at an interface between the putative S1-S2 interaction region, significantly weakens the binding of sS1 to proximal S2. H251N is next to R453C on the surface of the myosin mesa, also located at the interface between the putative S1-S2 interaction region, whereas D239N (in Switch-1) is remote from this region.

To test if H251N also disrupts the S1-S2 interaction, we used microscale thermophoresis (MST) to assess the relative binding affinities of WT and mutant sS1s with human proximal S2, as described in Nag et al. (*25*). MST measurements revealed, as for the R453C mutation (*25*), that there was a significant decrease in sS1-proximal S2 binding affinity due to the H251N mutation (200 ± 30 μM; 8 runs and 3 myosin preparations) in comparison to WT (45 ± 10 μM; 10 runs and 5 myosin preparations), but no change was observed for D239N (60 ± 8 μM; 5 runs and 2 myosin preparations) (**Figure 4b, Table 2**), which is located in a region where there is no proposed S1-S2 interaction.

**Figure 4.**
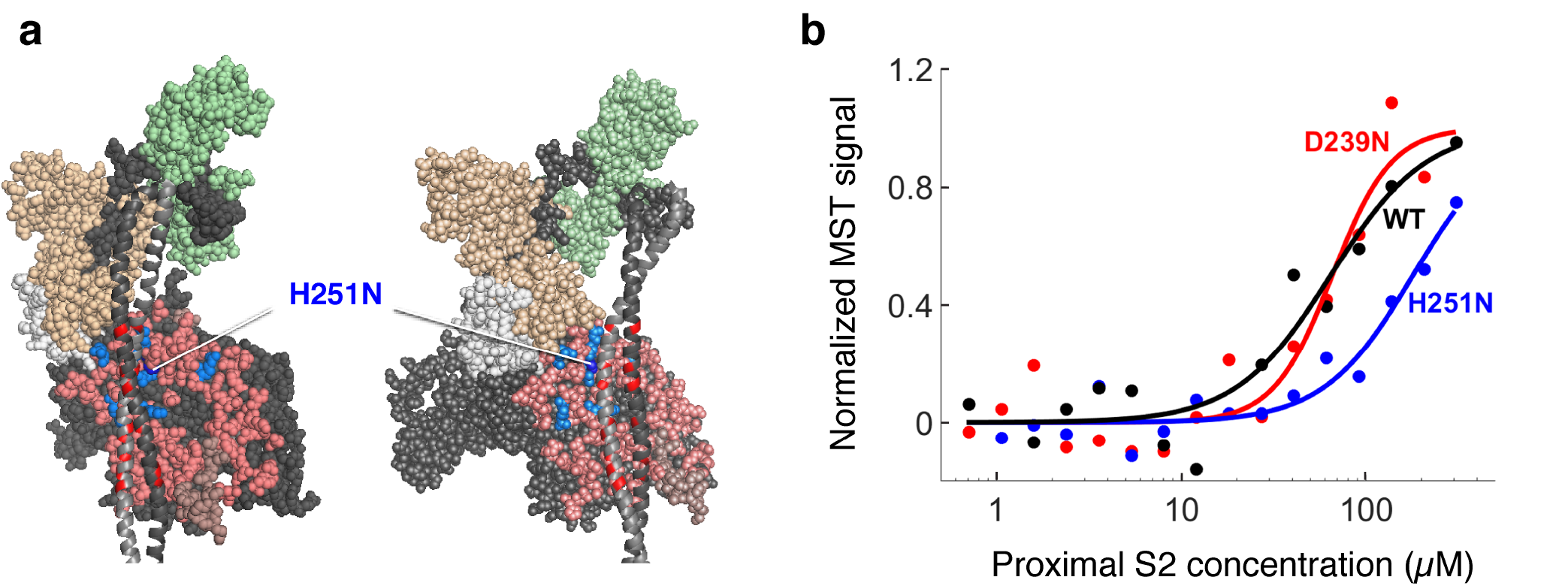
Binding of H251N and D239N human β-cardiac sS1s to human cardiac proximal S2. (**a**) Structural model of homology-modeled sequestered heads of human β-cardiac S1 based on the 3D-reconstructed structure of tarantula skeletal muscle myosin thick filaments by Alamo et al. (*44*). H251N is at the interaction site of S1 and S2, while D239N is not. (**b**) Representative MST binding experiments using human β-cardiac sS1 tagged with a C-terminal eGFP.

## Discussion and Summary

Elucidating the effect of HCM mutations on human β-cardiac myosin function and contractility at a molecular level has been challenging due to the small differences seen in the molecular parameters such as intrinsic force and duty ratio, some of which contribute to hypercontractility and others to hypo-contractility. Thus, how these mutations cause the typical hyper-contractile phenotype of clinical HCM is unclear.

Even though the incidence of pediatric HCM is low, the prognosis tends to be worse than for adult onset HCM, with a much higher incidence of morbidity and mortality (*4*). We predicted that mutations linked predominantly to an earlier onset and more severe clinical phenotype would show greater changes at the molecular level in comparison to previously studied HCM mutations that most often cause adult-onset disease. In this paper, we present the molecular characterization of two early-onset mutations, H251N and D239N, which show very large increases in function for all the human β-cardiac myosin biomechanical parameters measured. D239N is in the switch I region of the nucleotide binding pocket. Mutagenesis studies of residues in this region of myosin from a variety of sources have revealed their importance in coupling the energy of ATP hydrolysis to lever arm movement (*36, 37*) while structural studies suggest their importance in nucleotide release (*38, 39*). The velocity and ATPase rates of D239N are the highest of any of the human β-cardiac S1 mutations previously measured. H251 N is in the central β-sheet that undergoes twisting upon binding of ATP and communicates to the 50K domains to release actin. This allows switch II to move into the closed position and ATP to be hydrolyzed (*39*). While these two mutations are located in structurally different regions of the head, they both could alter how nucleotide binding translates to actin binding and force production. This idea is supported by the large changes observed in biomechanical properties.

In the case of H251N human β-cardiac myosin, an additional parameter that may be increased by the mutation is N_a_. The loaded motility assay does not account for the structural constraints of the assembled sarcomere and therefore does not account for N_a_. Thus, including this parameter to estimate the ensemble force may lead to greater differences in comparison between H251N and WT than is shown in Figure 3D.

To the best of our knowledge, H251N is the first HCM mutation to show increases in function for all measured parameters, and D239N shows increases for all parameters, except the number of available heads. Moreover, ensemble force studies carried out on these motors using the loaded in vitro motility assay show elevated velocities for both D239N and H251N human β-cardiac sS1 at all utrophin concentrations, thereby suggesting a larger power output (P = force x velocity) for the mutant myosins. Taken together, these data show increased function above what has been previously published for myosin HCM mutations, supporting the idea that, in concert with the more severe clinical phenotype, these early-onset mutations show more severe effects at the molecular level.

## Materials and methods

### Expression and purification of proteins

Recombinant human β-cardiac sS1 containing HCM causing mutations were co-expressed with a FLAG-tagged human essential light chain in C2C12 mouse myoblast cells using adenoviral vectors, and purified using FLAG affinity and ion exchange chromatography as described (*19*). The myosin construct includes either a C-terminal 8 amino acid peptide (RGSIDTWV) with, which acts as an affinity clamp (AC) that binds the PDZ-18 protein (myosin-AC), or a C-terminal GFP, which binds to an anti-GFP antibody (myosin-GFP).

C2C12 cells were grown at 37 °C and 8% CO_2_ in growth medium (DMEM + 10% fetal bovine serum + 1x pen-strep). 10 plates of confluent C2C12 cells were differentiated to myotubes by adding differentiation medium (DMEM + 2% horse serum + 1x pen-strep). The cells were differentiated for 2 days. Next, the cells were infected with adenoviruses carrying the myosin sS1 and myosin flag-ELC in growth media ½ (DMEM + 5% fetal bovine serum + 1x pen-strep). The cells were infected for 4 days and then harvested. The medium was removed from the cells, and they were washed with ice cold PBS. Then 1 ml of lysis buffer (20 mM imidazole, 100 mM NaCl, 4 mM MgCl2, 1 mM EGTA, 1 mM EDTA, 0.5% tween-20, 1 mM DTT, 3 mM ATP, 1 mM PMSF, 10% sucrose and Roche protease inhibitors) was added to each plate and the cells were harvested. The cells were lysed using a dounce, and then centrifuged at 23000 rpms for 20 minutes using a Ti60 ultracentrifuge rotor. The supernatant was then incubated with 1mL of anti-flag resin for 2 hours at 4 °C. This allowed the sS1-ELC complex to bind to the resin. The resin was then gently spun at 2000 rpms for 2 min. The supernatant was discarded, and then the resin was washed with wash buffer (20 mM imidazole, 150 mM NaCl, 5 mM MgCl2, 1 mM EGTA, 1 mM EDTA, 1 mM DTT, 3 mM ATP, 1 mM PMSF, 10% sucrose and Roche protease inhibitors) and incubated in wash buffer overnight with TEV protease at 4°C to cleave off the Flag-tag. The next day, the resin was spun and the supernatant containing the myosin was collected and further purified using anion exchange chromatography.

Actin was prepared from bovine cardiac muscle (identical in sequence to human cardiac actin), as previously described (*40*). PDZ-18 protein is a chimeric protein (erbin PDZ+fibronectin) engineered to bind the PDZ binding peptide (RGSIDTWV) with picomolar affinity (*41*). The His-tagged PDZ-18 is expressed using the Rosetta (DE3) bacterial expression system and purified using nickel affinity columns. His-tagged human proximal S2 myosin is expressed and purified in the same way as PDZ-18. His-tagged utrophin is also expressed and purified similarly, but requires an additional inclusion body purification step as described in Aksel et al. (*28*).

### Actin-Activated ATPase assays

Only freshly prepared myosin-eGFP was used for this assay. Once the myosin was purified, it was buffer exchanged into ATPase buffer (10 mM imidazole, 5 mM KCl, 1 mM DTT, 3 mM MgCl_2_) using amicon ultracell 10K filters. Myosin concentration was measured using eGFP absorbance. To prepare F-actin, G-actin was dialyzed extensively into ATPase buffer to remove any residual ATP. Actin concentration was then measured using absorbance at 290 nm using a spectrophotometer. The steady-state actin-activated ATPase activity of the WT and mutant human β-cardiac myosins was determined using a colorimetric assay to measure inorganic phosphate production at various time points from a solution containing myosin, ATP and increasing amounts of actin filaments (*42*). All measurements were made at 23 °C. Kinetic parameters (i.e. k_cat_) were extracted from the data by fitting to the Michaelis-Menten equation to determine maximal activity (*26*).

### Unloaded and loaded in vitro motility assays

We performed unloaded and loaded motility assays on WT and mutant motor domains (using myosin-AC). We have observed that there was very little difference in motility observed between freshly purified myosin and frozen myosin. Flow chambers were constructed from nitrocellulose-coated cover slips mounted on glass slides, and set up as follows: first the PDZ-18 protein (3 μM) was flowed in, followed by myosin (0.3-0.4 μM). After bovine actin labeled with phalloidin-tetramethylrhodamine (TMR) was added, movement was initiated by addition of ATP in the presence of oxygen scavenging and ATP regeneration systems (*31*). Actin filaments were detected under the microscope and at least 4 movies of 30 seconds were recorded for each condition. All measurements were made at 23°C. To remove motors that bind actin irreversibly, purified motor preparations were incubated with excess bovine skeletal actin on ice in the presence of ATP, then pelleted by ultracentrifugation (TLA 100; 95,000 rpm) at 4°C just prior to performing the assay.

For loaded motility assay, utrophin-AC was added along with myosin-AC at a concentration between 0.2-3.2 nM, and the motility was measured for each concentration of utrophin. The velocity data was analyzed using the FAST algorithm, as described previously (*28*). For the loaded motility assay, it was essential to ensure that the concentrations of WT and mutant myosins were the same. We measured the concentration of myosin using the Bradford-Assay (Bio Rad).

### Optical Tweezers Assay

Details of the experiment and data-analysis are described elsewhere (*18, 19*). Briefly, dead-heads of myosin were removed as described above, to increase the probability of stroking heads binding to actin. The nitrocellulose coverslip was sparsely coated with 1.5 μm silica beads (Bangs Laboratories, Inc.), which were then coated with anti-GFP antibody (Abcam). Next the chamber was passivated with BSA (1 mg ml^-1^) to prevent nonspecific sticking. The beads were sparsely coated with eGFP myosin (~200 pM). Next, individual biotinylated and TMR phalloidin labeled actin filaments were attached between trapped, neutravidin-coated polystyrene beads (Polysciences). When an individual myosin molecule binds to the actin and strokes, the force generated by the myosin molecule is measured (*19*).

### MST Assay

We used microscale thermophoresis to investigate the effect of the HCM mutations on the myosin S1-S2 interaction. We used freshly prepared myosin sS1-eGFP, and bacterially expressed proximal S2 myosin (839-968). The proximal S2 construct includes the first 126 amino acids of S2 and it begins 4 residues before the end of S1. Due to the low affinity between S1 and S2, the S2 myosin was concentrated to >300 μM to get a binding curve. Both proteins were dialyzed into MST assay buffer (10 mM Imidazole, 100 mM KCl, 1 mM EDTA, 2 mM MgCl_2_, 1 mM DTT, 500 μM ADP and 0.05% tween). Before using them for the assay, both proteins were centrifuged at 100,000 rpms (TLA 100 rotor) for 20 min to remove any aggregates. For the assay, we used 16 serial dilutions of S2 myosin starting at > 300 μM, with 2-fold dilution for every subsequent sample. The myosin sS1 was kept constant at 50 nM. The affinity measurements were performed using a Nanotemper thermophoresis set up at MST power = 60. The data was fit to the Hill equation. There were preparation-to-preparation differences in the binding affinity, and the WT K_D_ ranged between 35–50 μM; however, the relative difference between WT and mutant myosin was constant.

### Development of human β-cardiac myosin protein models

We developed human β-cardiac myosin S1 models based on the known motor domain structural data to best represent the human β-cardiac myosin, as described in Homburger et al (*23*). In brief, we retrieved the protein sequence of human β-cardiac myosin and the human cardiac light chains from UNIPROT database (*35*): myosin heavy chain motor domain (MYH7) - P12883, myosin essential light chain (MLC1) - P08590, and myosin regulatory light chain (MLC2) - P10916. We used a multi-template homology modeling approach to build the structural coordinates of MYH7 (residues 1-840), MCL1 (residues 1-195), MCL2 (residues 1–66) and S2 (residues 841-1280). We obtained the three dimensional structural model of S1 in the pre- and post-stroke states by integrating the known structural data from solved crystal structures, as described (*23*).

The S2 region is a long coiled-coil structure; hence we used the template from the Myosinome database (*20*). Modeling was done using the MODELLER package. Visualizations were performed using PyMOL version 1.7.4 (www.pymol.org).

## Acknowledgements

The authors would like to thank Dr. Leslie Leinwand and Dr. Robert Knight for personal communication about these mutations, Shirley Sutton for helping with virus preparation and cloning, Dr. Darshan Trivedi and Dr. Suman Nag for preparation of proximal S2, and for data acquisition and analysis of MST experiments. We also thank the rest of the Spudich Lab members for helpful discussions. The research was funded by NIH grants GM33289 and HL117138 (J.A.S.), Lucile Packard Child Health Research Institute Postdoctoral Fellowship (A.S.A, K.B.K), Stanford ChEM-H Postdocs at the Interface Award (A.S.A., K.B.K.), Stanford CVI Postdoctoral Award (A.S.A.), NIH T32 Training Grant (K.B.K), American Heart Association Postdoctoral Fellowship (A.S.A.), Stanford Spectrum Translational Medicine training grant (C.L.), Stanford CMB training grant (C.L.), and Stanford Bio-X fellowship (C.L.). J.A.S. is a founder of Cytokinetics and MyoKardia and a member of their advisory boards. K.M.R. is a member of the MyoKardia scientific advisory board.

## Author Contributions

Purified myosin and actin were prepared by A.S.A. with help from K.B.K. Unloaded and loaded *in vitro* motility were carried out by A.S.A., actin-activated ATPase by A.S.A. and K.B.K., intrinsic force measurements by C.L. and S.S.S., and MST experiments by A.S.A. Data analysis was done by A.S.A., K.B.K., and S.S.S. Molecular modeling was done by J.A.S. The manuscript was written and edited by A.S.A., K.B.K., D.B., K.M.R. and J.A.S.

